# MS Western, a Method of Multiplexed Absolute Protein Quantification is a Practical Alternative to Western Blotting

**DOI:** 10.1101/156943

**Authors:** Mukesh Kumar, Shai R. Joseph, Martina Augsburg, Aliona Bogdanova, David Drechsel, Nadine L. Vastenhouw, Frank Buchholz, Marc Gentzel, Andrej Shevchenko

## Abstract

Absolute quantification of proteins elucidates the molecular composition, regulation and dynamics of multiprotein assemblies and networks. Here we report on a method termed MS Western that accurately determines the molar abundance of dozens of user-selected proteins at the sub-femtomole level in whole cell or tissue lysates without metabolic or chemical labelling and without using specific antibodies. MS Western relies upon GeLC-MS/MS and quantifies proteins by *in-gel* co-digestion with an isotopically labelled QconCAT protein chimera composed of concatenated proteotypic peptides. It requires no purification of the chimera and relates the molar abundance of all proteotypic peptides to a single reference protein. In comparative experiments, MS Western outperformed immunofluorescence Western blotting by the protein detection specificity, linear dynamic range and sensitivity of protein quantification. To validate MS Western in an *in vivo* experiment, we quantified the molar content of zebrafish core histones H2A, H2B, H3 and H4 during ten stages of early embryogenesis. Accurate quantification (CV<10%) corroborated the anticipated histones equimolar stoichiometry and revealed an unexpected trend in their total abundance.

## INTRODUCTION

Despite well-known technical limitations and numerous application pitfalls, Western blotting (WB) remains one of the most widely used tools in analytical biochemistry (reviewed in (1–7)). WB conveniently provides a semi-quantitative estimate of the protein abundance directly from crude cell or tissue extracts. Quantification capabilities of WB, particularly its linear dynamic range, have been improved by using secondary antibodies bearing fluorescent labels and advanced systems for the optical readout of the abundance of recognized protein bands (8). This, however, has not alleviated the critical requirement of having antibodies with high and specific affinity towards target proteins (9).

Within the last two decades a variety of mass spectrometry based methods for targeted absolute protein quantification relying on isotopically labelled peptide / protein standards (*e.g*. AQUA (10), PSAQ (11), FLEXIQuant (12), prEST (13) or QconCAT (14, 15)); relative proteome-wide quantification using chemical or metabolic labelling *(e.g.* ICAT (16), TMT (17), iTRAQ (18, 19), SILAC (20), Super SILAC (21)), as well as label-free proteome quantification (22–26) have been developed. As an alternative to WB of SDS-extracted membrane proteins Arnott *et al* developed a method of SRM quantification of pre-selected pairs of ICAT-labeled peptides enriched by affinity chromatography prior to LC-MS/MS (27). However, these and other developments did not replace WB *en masse*, although it had been suggested that the field would strongly benefit from routine use of a targeted antibody-independent quantification of proteins by mass spectrometry (28).

Because of the attomole sensitivity, protein identification confidence, quantification accuracy, analyses throughput (reviewed in (29)) and, last but not least, the availability of high-end mass spectrometers proteomics has had a major impact on the entire field of molecular and cell biology. However, it is often perceived as a tool for monitoring global proteome-wide perturbations that is too cumbersome and inflexible for hypothesis-driven studies encompassing a limited selection of proteins that need to be quantified in many biological conditions. High costs and technical hurdles of proteome-wide labelling of tissues or entire model organisms with stable isotopes; cumbersome preparation of clean protein extracts; inconsistent quality of synthetic peptide standards; biased quantification of membrane and modified proteins are common bottlenecks in targeted proteomics applications.

Here we report on a method we termed MS Western that provides multiplexed absolute (in moles) antibody-free quantification of dozens of user-selected proteins from unlabelled cell and tissue lysates that combines sample preparation versatility of conventional WB with the specificity, accuracy and sensitivity of LC-MS/MS.

## EXPERIMENTAL PROCEDURES

### Chemicals and reagents

All reagents were of the analytical grade or better quality. LC-MS grade solvents were purchased from Fisher Scientific (Waltham, MA); formic acid (FA) from Merck (Darmstadt, Germany), Complete Ultra Protease Inhibitors from Roche (Mannheim, Germany); Trypsin Gold, mass spectrometry grade, from Promega (Madison, USA); restriction enzymes and buffers from New England BioLabs (Ipswich, MA); benzonase from Novagen (Gibbstown, NJ); other common chemicals and buffers were from Sigma-Aldrich (Munich, Germany). Pre-cast 4 to 20 % gradient 1mm thick polyacrylamide mini-gels were from Anamed Elektrophorese (Rodau, Germany).

### Protein standards and amino acids

Protein standards: bovine serum albumin (BSA), glycogen phosphorylase (GP), alcohol dehydrogenase (ADH), enolase (ENO) and ubiquitin (UBI) were purchased as a lyophilized powder from Sigma-Aldrich. Their purity was checked by 1D SDS PAGE and by amino acid analysis (Functional Genomics Centre Zurich, Switzerland). Ampoules of Pierce BSA standard and of recombinant human histones were purchased from Thermo Fisher Scientific (Waltham, MA) and from New England BioLabs (Ipswich, MA), respectively. Amino acids were from AppliChem (Darmstadt, Germany); isotopically labelled ^13^C_6_^15^N_4_-L-arginine and ^13^C_6_-L-lysine were from Silantes (Munich, Germany).

### Protein analysis by GeLC-MS/MS

To visualize protein lanes gels were stained with Coomassie CBB R250 and gel slabs covering the mass range of *ca* +/−20 kDa off the Mr of target proteins were excised. Proteins were *in-gel* digested with trypsin (30) and recovered tryptic peptides were analyzed by nanoflow LC-MS/MS (31); full details on the GeLC-MS/MS procedure are provided in Supplementary Methods. Mass spectra were acquired in data-dependent acquisition (DDA) mode on LTQ Orbitrap Velos or Q Exactive HF mass spectrometers, both from Thermo Fisher Scientific (Bremen, Germany). DDA settings are provided in Supplementary Table S5.

### Data processing

To match peptides to target proteins, MS/MS spectra were searched by Mascot v.2.2.04 software (Matrix Science, London, UK) against a customized database containing sequences of all target proteins, human keratins and porcine trypsin (in total, 234 protein entries). We applied precursor mass tolerance of 5 ppm; fragment mass tolerance of 0.6 Da and 0.03Da for LTQ Orbitrap Velos and Q Exactive HF instruments, respectively; fixed modification: carbamidomethyl (C); variable modifications: acetyl (protein N-terminus), oxidation (M); labels: 13C(6) (K) and 13C(6)15N(4) (R); cleavage specificity: trypsin, with up to 2 missed cleavages allowed. Peptides having the ions score above 20 were accepted (significance threshold *p* < 0.05) and the quantification was only carried out if both light and heavy forms of the same peptide were identified by MS/MS and retention time of their XIC peaks matched. Xcalibur (Thermo Fisher Scientific) and Progenesis LC-MS v.4.1 (Nonlinear Dynamics, UK) software were used for extracting peptide features from LC-MS/MS datasets.

### Experimental design and statistical rationale

For benchmarking and validation of MS Western we designed four QconCAT chimeric proteins (CP01 to CP04) having MW in the range of 35 to 264 kDa and comprising proteotypic peptides from proteins of different properties (*e.g*. cytosolic, transmembrane), size (from 8 to 2065 kDa) and organismal origin. Knock-down experiments in HeLa cells were performed in biological triplicates and analyzed by LC-MS/MS in technical duplicates. MS Western benchmarking experiments were performed in technical duplicates for each of 15 samples made by successive dilution of a total protein extract of HeLa cells. Core histones were quantified in zebrafish embryos in three biological replicates and LC-MS/MS run were acquired in technical duplicates. Wherever applicable, we provide standard deviation (±SD), coefficient of variance (CV) and robust coefficient of variance (rCV) calculated as 1.4826 times the median absolute deviation (32).

### Quantification of proteins spiked into an *E.coli* lysate

The overnight culture (OD_600_=1.5) of *E.coli* strain BL21 (DE3) T1 pRARE was pelleted by centrifugation at 5000 rpm for 5 minutes at 4°C (JLA 8.1000 centrifuge from Beckman Coulter, Brea CA). The cell pellet was re-suspended in 2x PBS, lysed in the presence of protease inhibitors and the total protein content was quantified using Pierce BCA Protein Assay Kit (Thermo Fisher Scientific). Equimolar mixtures containing 25 fmol to 25 pmol of each protein standard (stock concentrations were quantified by amino acid analysis) were spiked into 50 µg of an *E.coli* protein extract and subjected to 1D SDS PAGE as above. Upon electrophoresis gel slices corresponding to the apparent MWs of spiked proteins were excised, co-digested with gel bands of CP01 and recovered peptides quantified by LC-MS/MS.

### Absolute quantification of human recombinant histone H4

Aliquots of a stock solution of recombinant human H4 with the concentration of 1mg/ml were appropriately diluted and mixed with equal volumes of 2x Laemmli buffer. Three samples containing 0.05, 0.1 and 0.3 µg of histone H4 were subjected to 1D SDS PAGE as described above. In parallel, the CP02 and an aliquot containing 0.066 µg of Pierce BSA standard were run on a separate gel. The excised bands of histone H4, CP02 and BSA were mixed and co-digested with trypsin. After *in-gel* co-digestion and extraction, tryptic peptides were reconstituted in 46 µL of 5% aqueous FA and 5 µL were analyzed by LC-MS/MS.

### Knock-down experiments in HeLa cells

AKT1, CAT, PLK1, and TUBA4A genes were knock-down (KD) in HeLa cells using RNAi (Eupheria Biotech, Dresden, Germany). KD experiments were performed in triplicate; renilla luciferase transfections as a RNAi specificity control were performed on a separate plate. Further details are provided in Supplementary Methods; details on antibodies and RNAi probes are in Supplementary Tables S6 and S7, respectively.

### Comparison of protein quantification by MS Western and LI-COR Odyssey

HeLa cells were cultured in Dulbecco’s modified Eagle’s medium supplemented with 10% fetal calf serum and 1% penicillin-streptomycin (Gibco^TM^ Life Technologies). 1×10^7^ HeLa cells were resuspended in 100 µL RIPA buffer containing Complete Protease Inhibitor Cocktail, EDTA-free (Roche). Equal volume of 2x Laemmli buffer was added and incubated at 80°C for 15 minutes with intermittent vortexing. The supernatant was transferred to a new vial and further diluted with RIPA / 2x Laemmli buffer (1:1, v/v). Samples obtained at each dilution step were subjected to 1D SDS PAGE on two different gels. One gel was analyzed by LI-COR Odyssey and another by MS Western using bands of CP03 and BSA as standard and reference proteins, respectively. In each dilution step an equivalent amount of target proteins was digested and injected into LC-MS/MS or imaged by the Odyssey.

### Monitoring the kinetics of *in-gel* digestion of HeLa proteins

Protein extracts of *ca*. 1× 10^7^ HeLa cells were prepared as described above, however in one series of experiments the same amount of cells was homogenized in a twice larger volume of the buffer. Aliquots of cell extracts each equivalent to 4% of the total amount of recovered protein material were loaded onto multiple lanes of polyacrylamide mini-gels. Upon SDS PAGE, gel slabs whose Mr corresponded to TUBA and CAT proteins were excised; gel bands of CP03 and of the reference protein BSA were mixed with each gel slab and all samples were in parallel digested with trypsin. After the specified periods of time one sample per each digestion experiment was withdrawn, peptides were extracted from the entire in-*gel* digest and quantified by LC-MS/MS as described above. Each sample was analysed in technical duplicates. The amount of protein digested at each time point was calculated by averaging the amounts of five independently quantified peptides from TUBA and BSA and of three peptides from CAT.

### Absolute quantification of histones in zebrafish embryos by MS Western

Wild-type (TLAB) zebrafish embryos were dechorionated immediately upon fertilization, synchronized and allowed to develop to the desired stage at 28°C. Ten embryos per developmental stage (except five embryos for 1-cell stage) were manually deyolked and snap frozen in liquid nitrogen. Samples were boiled in the Laemmli buffer at 98°C for 10 minutes and subjected to 1D SDS PAGE. A single gel slab containing histones was excised from each sample lane and histones were quantified by MS Western using bands of CP02 and BSA as standard and reference proteins, respectively. 10% of the total amount of recovered tryptic peptides were injected into LC-MS/MS.

### Absolute quantification of histones in zebrafish embryos by LI-COR Odyssey

Zebrafish embryos were collected at the specified developmental stages (*n* = 5 for H3 and H2B; *n*=10 for H2A and H4). Embryos were processed as described above and total protein extracts were subjected to 1D SDS PAGE. Proteins were blotted onto a nitrocellulose membrane (GE Life Sciences). Primary antibodies (Supplementary Table S8) were incubated at room temperature for 1 hour or at 4°C overnight; secondary antibodies (Supplementary Table S8) were incubated at room temperature for 45 min. Proteins were quantified by LI-COR Odyssey using tubulin as a loading control. Standards of recombinant human histones were used for making calibration plots for the quantification of corresponding zebrafish homologues.

### Data deposition

Proteomics data have been deposited at the ProteomeXchange Consortium *via* the PRIDE (33) partner repository with the dataset identifier PXD005654 and doi: 10.6019/PXD005654. Temporary login for reviewers is provided under username: and password: HgrBxHbm.

## RESULTS AND DISCUSSION

### MS Western: protein quantification concept and workflow

Effectively, MS Western merges the three established analytical approaches: GeLC-MS/MS (30, 31, 34–36); proteotypic peptides clubbed with “top *N* peptides” protein quantification (31, 37–42) and QconCAT synthesis of isotopically labelled protein chimeras comprising concatenated sequences of proteotypic peptides (14, 15, 32). We termed this method as MS Western to underscore that it is targeted (rather than global), quantitative, relies on SDS PAGE of crude protein extracts and in this way is in line with classical Western. However, because of mass spectrometry readout, it requires no blotting and, most importantly, no antibodies.

To quantify a protein by MS Western we first selected a few (typically, three to six) proteotypic peptides (37, 39) in preliminary GeLC-MS/MS experiments, which also verified the position of bands of target proteins at the electrophoresis lanes separating crude protein extracts. However, peptides could be also picked from collections of LC-MS/MS spectra (43, 44) or predicted by software (45, 46).

In the same way, we further selected proteotypic peptides from two reference proteins - in this work we used glycogen phosphorylase (GP) and bovine serum albumin (BSA). Peptide sequences from the target and reference proteins were concatenated *in-silico* in an arbitrary order except that peptides from the same protein were positioned successively. The entire stretch of peptide sequences was flanked at the N- and C-termini with the sequences of twin-strep-tag followed by a 3C protease cleavage site and His-tag, respectively (Fig. 1). These tags protect target peptides from exopeptidase degradation and, only if deemed necessary, could be used to enrich the expressed chimera from a whole cell lysate. Altogether we designed four project-specific chimera proteins (CP) ranging in size from 35 to 264 kDa that encoded, in total, more than 300 proteotypic peptides from 58 individual proteins (Supplementary Fig. S1, S3, S5 and S7). The design rationale was the same as in QconCAT proteins (14, 15) and here we use CP acronym solely for the presentation clarity. All CPs made in this work were highly expressed (Supplementary Fig. S2, S4, S6 and S8) and incorporated more than 99.5% of heavy arginine and lysine residues (Supplementary Fig. S9).

**Fig. 1.**
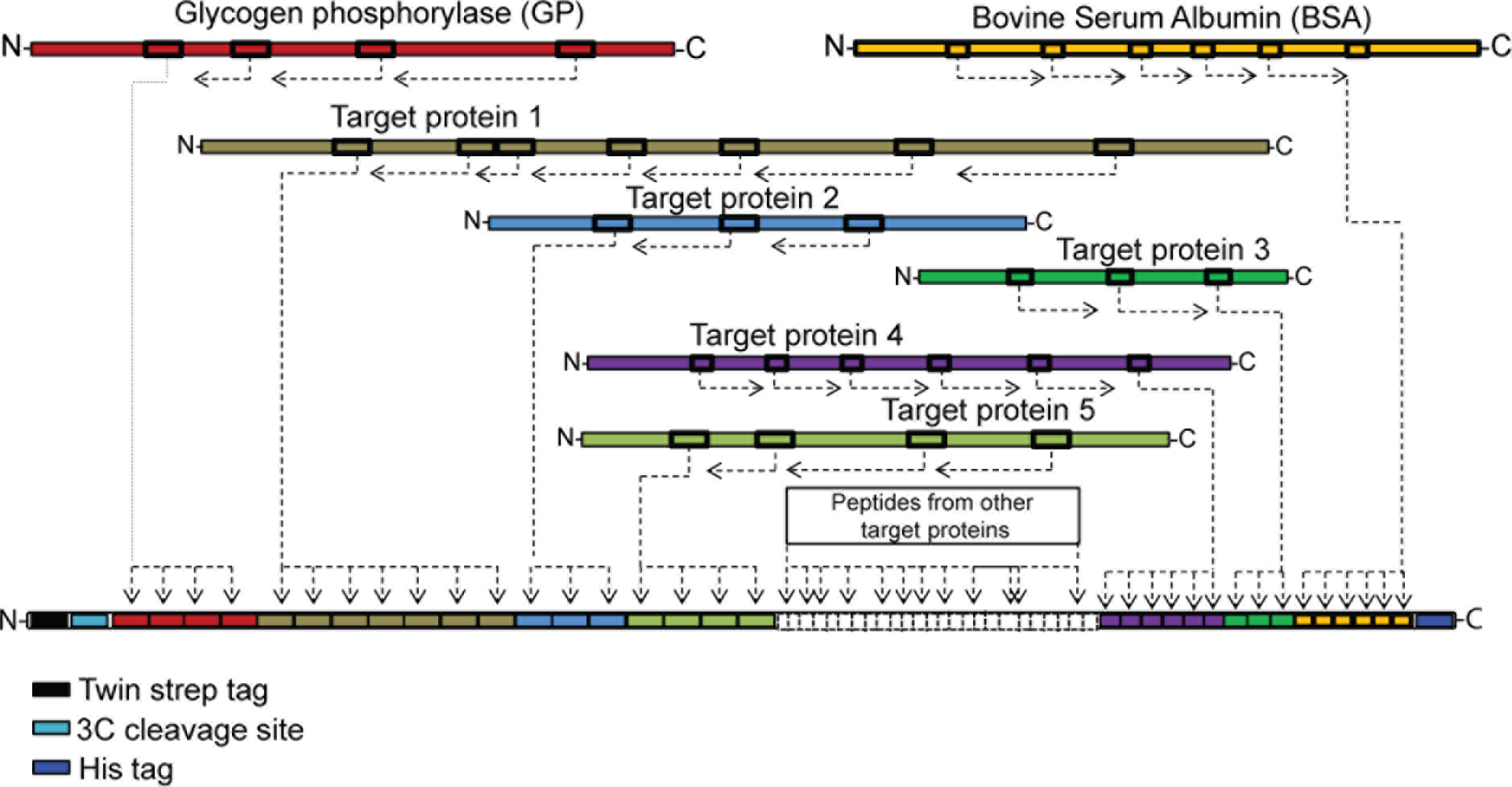
Modular organization of chimera proteins (CP) used in MS Western‥. Sequences of proteotypic peptides (schematically shown as boxes) from each target protein (colour-coded) are in-silico concatenated into a single chimera, flanked with peptide sequences from the reference proteins GP (at the N-terminus side) and BSA (at the C-terminus side) and with two affinity tags together with 3C cleavage site.

In contrast to the relative (fold change) quantification, absolute (in moles) quantification critically depends on the exactly known concentration of internal standard(s). In QconCAT and related methods it is usually determined by the amino acid analysis or photo- or colorimetric assays (15). This, however, requires highly purified CPs and is prone to batch-to-batch variations and interlaboratory inconsistency. Instead, MS Western utilizes a simple workaround solution that requires no CP purification (Fig. 2).

**Fig. 2.**
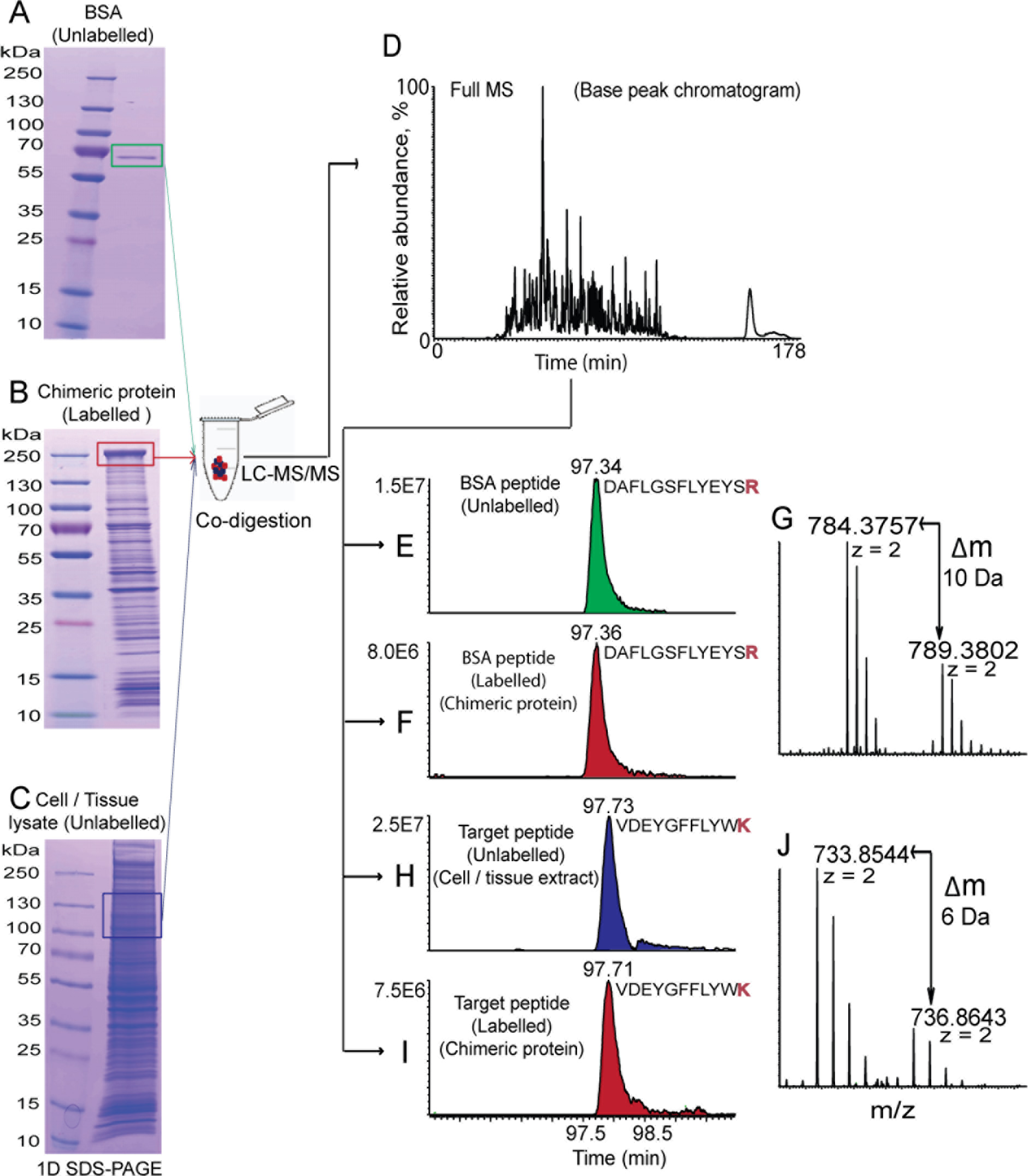
Workflow of the absolute protein quantification by MS Western. Gel band containing the known amount of the reference protein BSA (**A**); gel band of CP metabolically labelled with ^15^N_4_, ^13^C_6_-Arg and ^13^C_6_-Lys (**B**) and the gel slab excised at the range of expected Mr of the target protein(s) (**C**) are co-digested with trypsin and recovered peptides are analyzed by LC-MS/MS (**D**). The CP is quantified by comparing the abundances of XIC peaks of unlabelled (from BSA reference) (**E**) and labelled (from the CP) (**F**) peptides (**G**). Next, the amount of target protein is inferred from the ratio of abundances of XIC peaks of matching unlabelled (from the target protein) (**H**) and labelled (from the CP) (**I**) proteotypic peptides (**J**) and from the amount of CP. For clarity, only two (out of several) matching pairs of labelled / unlabelled peptides are shown. A solubilised cell pellet in panel **B** was subjected to SDS PAGE without prior enrichment of the CP. Quantification of a peptide from norpA protein from D.*melanogaster* using CP04 chimera is shown as an example.

To quantify proteins of interest a cell or tissue lysate was subjected to 1D SDS PAGE. Gel slab(s) approximately matching the Mr of target protein(s) were excised and mixed with the band of the CP excised from a lane with *E.coli* lysate and with the band of a reference protein (*e.g*. BSA). The three bands were co-digested with trypsin, recovered peptides were analyzed by LC-MS/MS and quantified by considering the abundance of XIC peaks of matching pairs of light and heavy peptides. Molar abundance of each quantified protein (including CP) was calculated by averaging molar abundances of several (typically, three to six) proteotypic peptides. Gel bands of CP, BSA and the gel slab with target protein(s) were *in-gel* digested together, light and heavy peptides were extracted from the same digest and quantified in the same LC-MS/MS experiment. Hence, in MS Western workflow the molar amount of each target protein was inferred from the molar amount of the CP, which in turn was referenced to the known molar amount of the BSA standard. However, there was no need in purifying CPs to homogeneity or maintaining their stock solutions with the exactly known concentration. In all experiments absolute (molar) quantities of all CPs were referenced to the same commercial protein standard (Pierce BSA) with the guaranteed chemical purity, stock concentration and batch-to-batch reproducibility.

MS Western protocol should be adjusted according to the Mr and abundance of target proteins. Coomassie staining was not intended to visualize bands of target proteins. Instead, proteins were excised within slightly larger (typically, +/− 20 kDa) gel slabs (Figure 2C) centred at their expected Mr and verified by preliminary GeLC-MS/MS. If a target protein smeares across several gel slices, molar amount (but not raw peak areas) of each peptide in each slice should be summed up. For optimal quantification the abundance ratio between light and heavy peptides should be adjusted by considering the isotopic purity (≥99.5%) of heavy arginine and lysine amino acids, rather than large (better than 10^4^-fold) linear dynamic range of Orbitrap analyzers (47). We typically co-digested *ca*. 100 fmols of CPs, depending on the abundance of target proteins. If required, the detection sensitivity could be further enhanced by using DDA with an inclusion list of expected *m/z* of target peptide precursors, by t-SIM or parallel reaction monitoring (PRM) (48).

Taken together, a vastly simplified MS Western workflow offers several experimental advantages. Firstly, both CP and target proteins are independently quantified in the same experiment by several proteotypic peptides and it is expected that they should produce quantitatively consistent determinations (31, 40). Secondly, MS Western protocol only requires femtomole amounts of isotopically labelled standards. All four CPs were produced with high yield (Supplementary Fig. S2A, S4A, S6A and S8A) and *ca* 1 mL of the cell culture provided them in hundred picomole quantities. In our experience none of expression experiments failed or low yield of CPs required their enrichment prior to MS Western quantification. Also, we did not maintain pre-quantified CP stocks, but simply excised their bands directly from a gel loaded with *E.coli* lysate. Thirdly, since proteins were solubilized in 4% SDS and analyzed by GeLC-MS/MS the quantification was not affected by the solubility or purity of CPs and target proteins. SDS PAGE also improved both the dynamic range and sensitivity of peptide detection. In contrast to a gel-free “mass western” approach by Arnott *et al* (27) that is also compatible with SDS extraction of proteins our method enables their absolute (rather than relative) quantification (also including membrane proteins) and uses no chemical derivatization and /or multistep clean-up of heavy and light pairs of proteotypic peptides. And finally, in contrast to conventional WB, MS Western does not rely on antibodies and ordering a synthetic gene at the current price level of *ca* 10 to 15 Eur per peptide with guaranteed quality and a few week delivery time is simply incomparable with costs and efforts required for generating and validation of reliable monoclonal antibodies.

### Validation and properties of MS Western

Although the concept of MS Western looked appealing, a major question remained if *in-gel* codigestion of relatively pure CPs, of target protein(s) embedded into a polyacrylamide gel matrix together with abundant background proteins and of pure BSA standard delivered the same yield of proteotypic peptides?

To answer this question, we designed and expressed *ca*. 42 kDa chimera CP01 (Supplementary Fig. S1 and S2A) comprising 31 proteotypic peptides from five commercially available protein standards: GP, BSA, ENO, ADH and UBI (Supplementary Table S1). GeLC-MS/MS of its gel band yielded 100% sequence coverage (Supplementary Fig. S2B) and confirmed ≥99.5% isotopic labelling efficiency (Supplementary Fig. S9).

Gel bands of each of these five target proteins were co-digested with the band of CP01. The extracted peptides were analyzed by LC-MS/MS and relative abundances of light (normalized to all light) and heavy (normalized to all heavy) proteotypic peptides were compared (Supplementary Fig. S10). We observed that within 31 peptide pairs the relative abundances varied by less than 5% (Supplementary Fig. S10), despite light and heavy peptides originated from structurally different chimera and endogenous proteins. Indeed, prior to *in-gel* digestion chimera, target and reference proteins were fully denatured first by SDS and then by acetic acid / methanol during Coomassie staining. Also, because of pre-separation of crude extracts by SDS PAGE, only a small fraction of a background proteome was co-digested together with target protein(s). In line with previous findings (32), we observed no noticeable impact of the size and composition of both target protein and CPs and speculated that SDS PAGE and *in-gel* digestion might even relax the constraints (49) applied for the selection of proteotypic peptides.

In only a few instances (Supplementary Fig. S10B and S11F) not all relative abundances matched because of trypsin miscleavages or post-translational modifications (PTMs). However, irrespectively of why they mismatched, “problematic” pairs of peptides could be spotted by their discordant relative abundances and disregarded from the protein quantification.

We next asked if the likeness of peptide relative abundances in *in-gel* digests of CP and of target proteins warranted their accurate molar quantification. As a test bed, we used the CP02 comprising peptides from the four core histones H2A, H2B, H3 and H4 from *D.rerio* (Supplementary Fig. S3, S4 and Supplementary Table S2). We also obtained a standard of human recombinant histone H4 supplied as a stock solution with the exactly known concentration. Human and zebrafish histones H4 share 99% full-length sequence identity (Supplementary Fig. S12) and we tested if human H4 could be quantified using identical peptides from CP02 whose molar abundance was referenced to the BSA standard. Three aliquots containing different amounts of human H4 were subjected to SDS PAGE and quantified by MS Western. Relative abundances of light peptides from BSA and histone H4 and corresponding heavy peptides from CP02 were in a good agreement (Fig.3A and Fig.3B, respectively). Relative abundances of one BSA (QTALVELLK) and one histone H4 (TVTAMDVVYALK) peptides mismatched because of trypsin miscleavage caused by the presence of flanking dibasic amino acid residues in the sequences of endogenous proteins (45, 50–52). When the areas of XIC peaks of miscleaved peptides were added into the calculation, the expected relative abundances were restored. Next for each peptide and each amount of histone H4 we calculated the ratio of relative abundances of their light (normalized to all light) and heavy (normalized to all heavy) peptide forms. If these normalized relative abundances remain the same then their ratio should be close to the value of 1.0, which was also consistent with our findings (Fig.3C). Altogether in the three independent experiments MS Western quantification relying on five proteotypic peptides (Fig.3B) correctly determined different molar amounts of histone H4 loaded on the gel (Fig.3D).

**Fig. 3.**
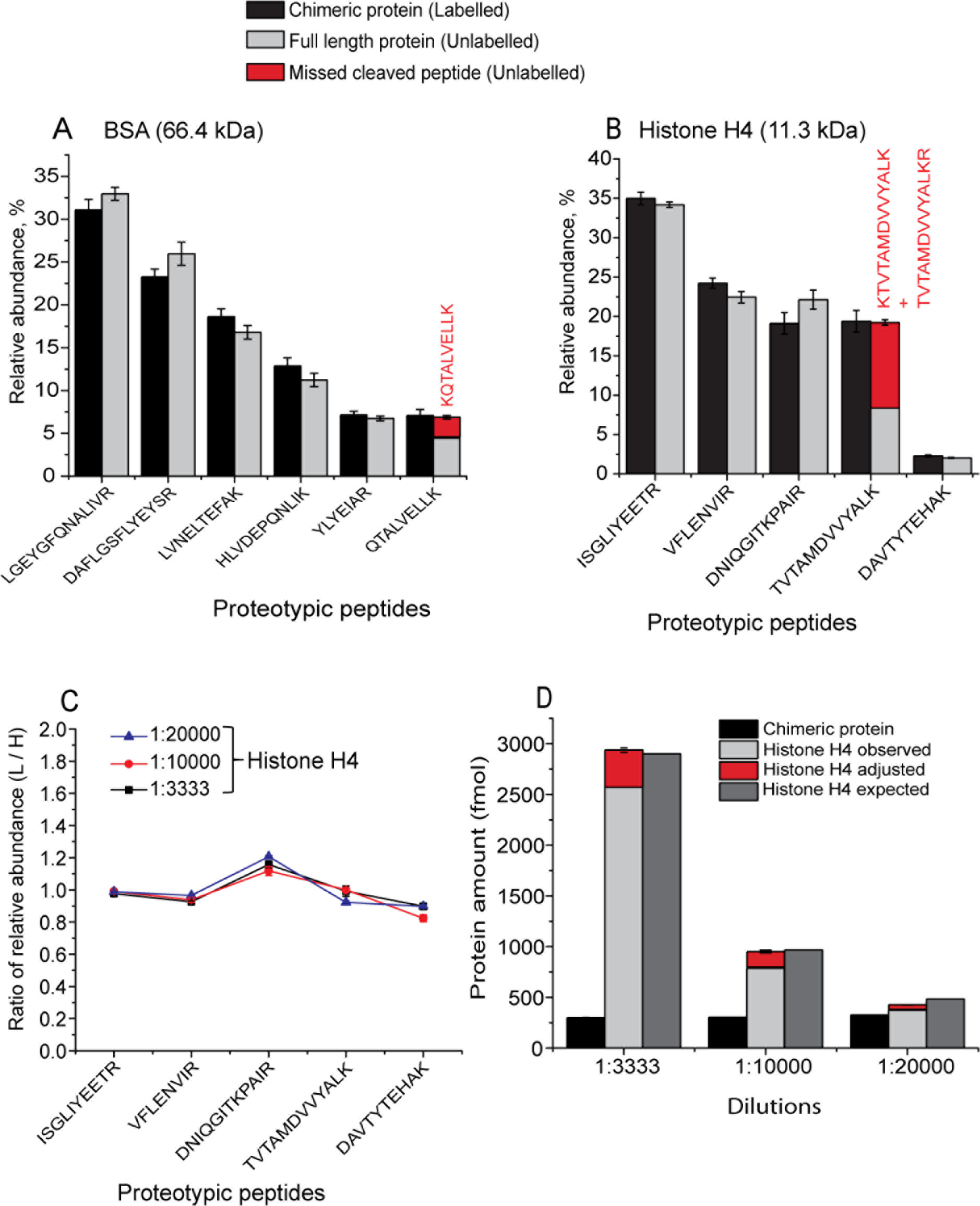
Validation of MS Western by absolute quantification of recombinant human histone H4. (**A**) Relative abundances of light and heavy BSA peptides produced by co-digestion of the bands of BSA and CP02, respectively. (**B**) Relative abundances of light and heavy peptides produced by co-digestion of the bands of recombinant human H4 and CP02, respectively. Abundances and sequences of miscleaved peptides complementing corresponding proteotypic peptides are in red. (**C**) Ratio of relative abundances of light and heavy peptides (as in panel **B**) at three different loadings of histone H4. (**D**) Loaded and MS Western quantified molar amounts of histone H4. The amount of co-digested CP02 is provided as a reference. Data are presented as mean ±SD (panels A and B *n*=3; panels C and D *n*=2).

Finally, we checked if the target, chimera and reference proteins were *in-gel* co-digested each at its own kinetics and if digestion rate matching was required for accurate absolute quantification? To this end, we monitored *in-gel* digestion of α-tubulin (TUBA, 50 kDa) and catalase (CAT, 60 kDa) in a SDS PAGE separated total protein extract from HeLa cells. Gel slabs were excised at the correspondent Mr, co-digested with bands of 72 kDa chimera protein CP03 (Supplementary Fig. S5, S6 and Supplementary Table S3) comprising proteotypic peptides from TUBA and CAT and with bands of the reference protein BSA. Peptides extracted from *in-gel* digests were quantified by LC-MS/MS to produce the kinetic plots in Fig. 4A, B and C. In the HeLa extract the abundance of TUBA and CAT differed by *ca* 100-fold (Fig. 4B, C) and they were digested together with *ca* 1100 comigrated background proteins. As a consistency check, we digested two loadings of the same extract whose total protein amount differed by 2-fold. In line with the previous report on *in-solution* digestion of chimera proteins (53), CP03 was digested *in-gel* within a few minutes. The digestion of TUBA and CAT was complete after *ca* 6 hr, while MS Western protocol relies on overnight digestion. Interestingly, not all proteotypic peptides were produced at the same rate (Fig. 4D, E). The ratio of relative abundances of light and heavy forms of most peptides plateaued at the expected value of 1.0 already after 3 hr, consistently with the kinetic plots in Fig. 4B, C. However, two peptides showed deviating trends. The ratio of relative abundances of peptide QTALVELLK was lower than expected (Fig. 4D, Supplementary Figure S13) because of miscleavage of native BSA (see also Fig. 3A). Consistently with earlier reports on the specificity of trypsinolisys of peptides flanked with successive Arg, Lys residues (45, 50, 51, 54) the yield of QTALVELLK did not improve at extended digestion times, while its release from CP03 was rapid and complete. Peptide QLFHPEQLITGK from TUBA was also produced at the slow rate, however its release was complete after 12 hours (Fig. 4E, Supplementary Figure S14). Importantly, biased yield of both peptides at all digestion time points was clearly reflected by deviating ratios of relative abundances of their light and heavy forms and supported the informed decision on accepting or excluding them from protein quantification.

**Fig. 4.**
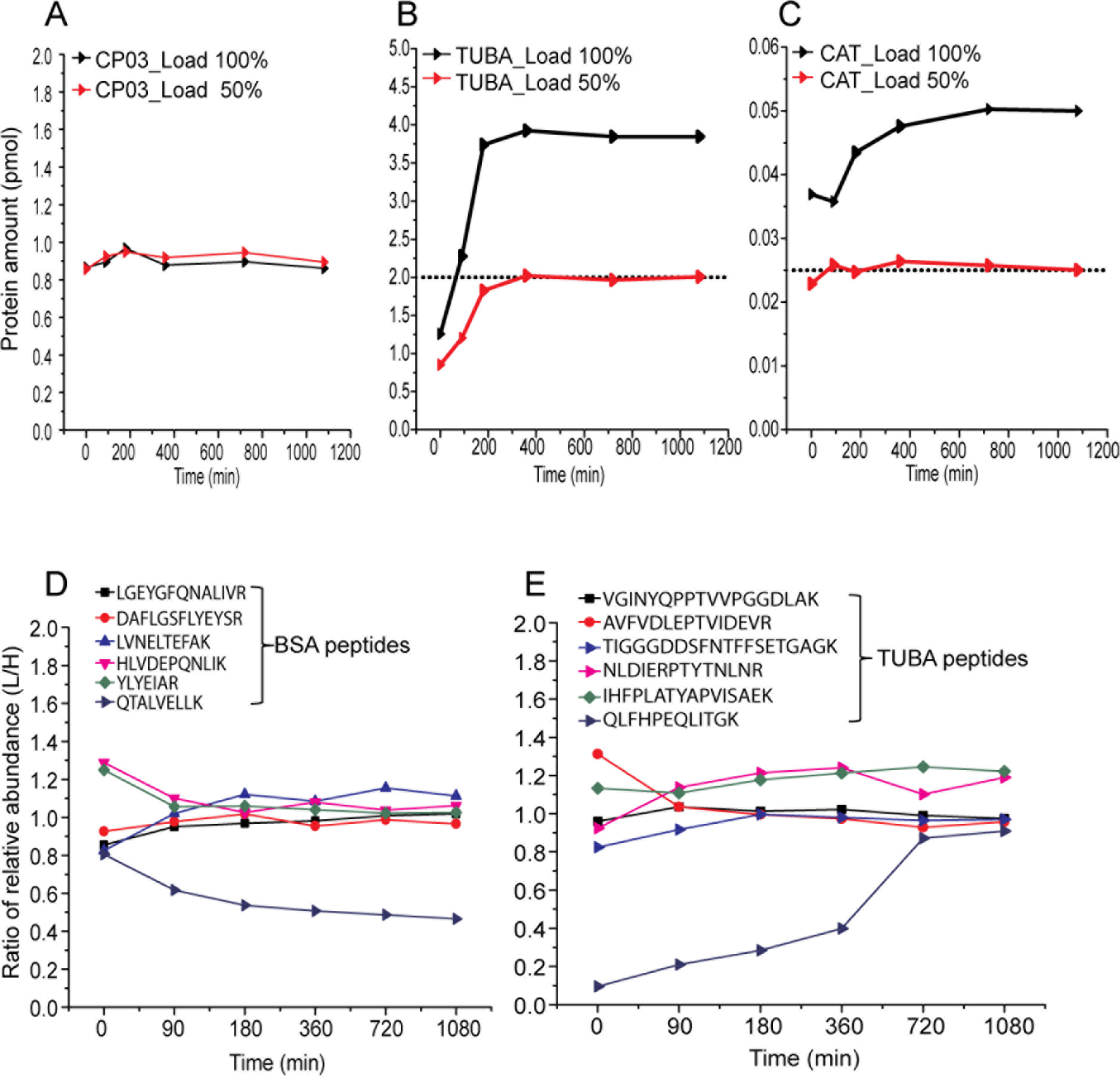
Fig. 4 Kinetics of n *in-gel* co-digestion of endogenous and chimeric proteins. (*A*) Digestion of CP03. (**B**) and (**C**) Digestion of TUBA and CAT, respectively. Dotted line indicates the expected 50% amount of proteins in the experiments with 2-fold diluted extract. (**D**),(**E**) Ratios of relative abundances of light and heavy peptides from BSA (reference protein) and TUBA, respectively. Plots designated as Load 100% and Load 50% stand for the digestion of proteins from an undiluted and 2-fold diluted HeLa extract, respectively. The amount of CP03 was the same in all co-digestions. Protein amounts at the y-axes indicate the average values of two technical replicas.

### Benchmarking protein quantification by MS Western

We further benchmarked MS Western quantification in three ways. First, we checked if its linear dynamic range was affected by a concomitant protein background. Second, we tested if both WB and MS Western produced quantitatively concordant determinations of endogenous proteins in whole cell lysates. Finally, we benchmarked MS Western sensitivity and dynamic range against currently the most advanced quantitative Western blotting system LI-COR Odyssey (further termed as Odyssey) that relies upon near-infrared immunofluorescent readout of the abundance of antibodies recognizing target proteins.

We first analyzed a series of eleven samples containing 25 fmol to 25 pmol of an equimolar mixture of five proteins (GP, BSA, ENO, ADH and UBI) spiked into 50 µg of *E.coli* protein extract. Each sample was subjected to SDS PAGE and the five proteins were quantified by MS Western by co-digesting their bands with the band of CP01. Closely migrating bands of ENO and ADH (ΔMr *ca* 10 kDa) were excised and digested together. Calibration graphs were produced by plotting the average of ratios of absolute abundances of light and heavy forms of proteotypic peptides pertinent to each protein against the injected amounts of protein digests and were linear within the entire range of protein loadings (Fig.5A). Calibration plots for 21 individual peptides used for making Fig.5A are provided in Supplementary Fig. S15 and suggested that peptides recovery was independent of protein size and percentage of acrylamide in digested gel slabs and was not affected by protein background.

**Fig. 5.**
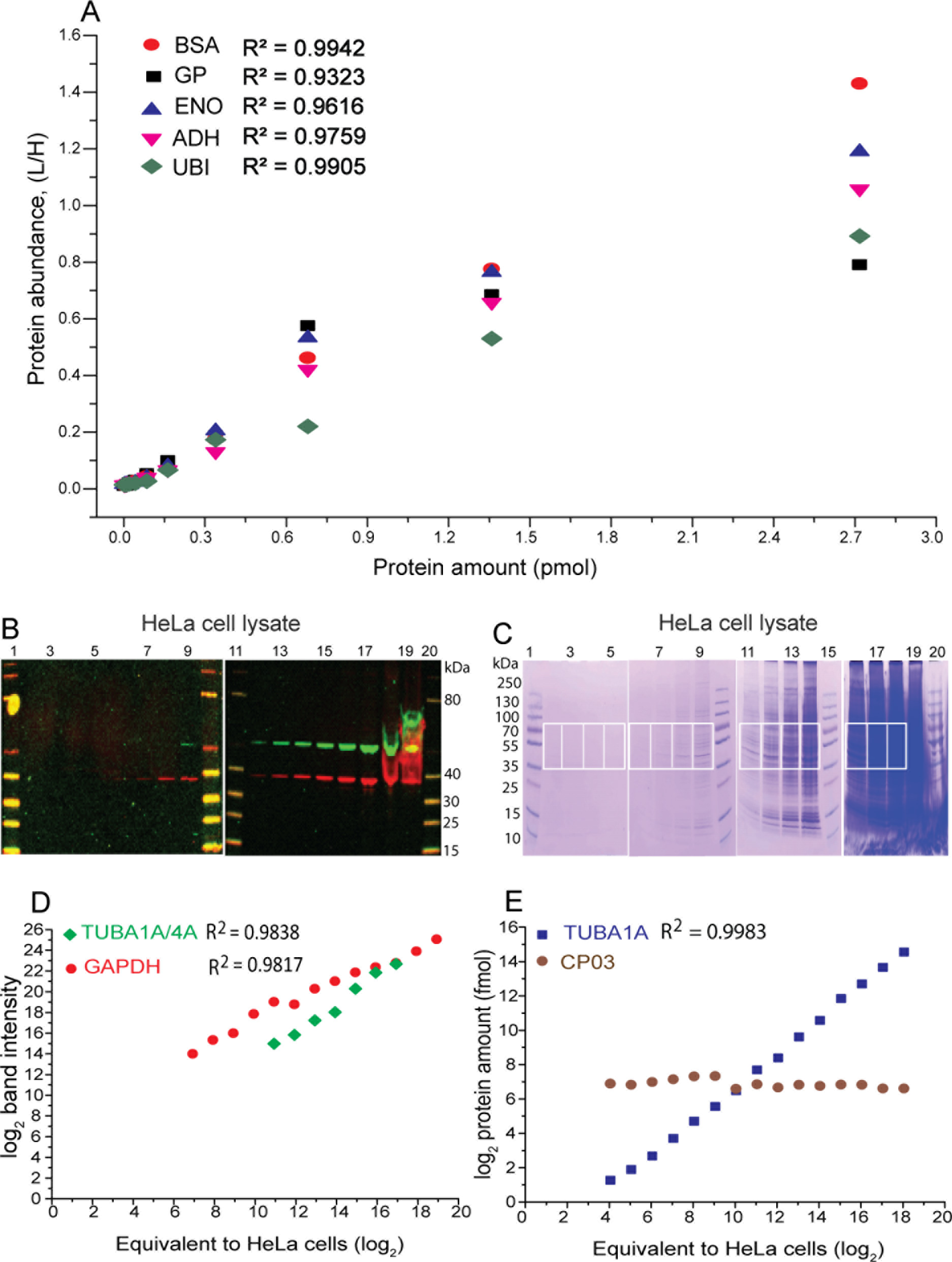
Benchmarking the protein quantification by MS Western. (**A**) MS Western quantification of proteins spiked into the constant background of *E.coli* lysate; y-axes: protein abundance expressed as an averaged ratio of abundances of light and heavy proteotypic peptides; x-axes: loaded protein amount, pmols (**B**) LI-COR Odyssey image of gels loaded with a successively diluted protein extract from HeLa cells. Bands with green and red fluorescence are TUBA4A/1A and GAPDH (loading control), respectively. (**C**) Coomassie stained gels with the same dilution series. Gel slabs excised from each lane for MS Western quantification of TUBA1A/4A are boxed. (**D**) Abundances of TUBA1A/4A from lanes 9,12,13,14,15,16,17 and GAPDH from lanes 5,6,7,8,9,12,13,14,15,16,17,18, 19 from the gel image in panel B as reported by the Odyssey. (**E**) Molar amount of TUBA1A and CP03 quantified by MS Western from slabs excised from lanes 2,3,4,5,6,7,8,9,11,12,13,14,16,17, 18 from the gel in panel **C**. Lanes 1,10,11,20 and lanes 1, 10, 15 and 20 are MW markers in panels **B** and **C**, respectively.

Next we tested if WB on LI-COR Odyssey and MS Western could provide concordant quantification of endogenous proteins in whole cell lysates. In contrast to MS Western, pure protein standards are required for the absolute quantification by the Odyssey because slopes of calibration plots relying on integral intensities of visualized bands differ between individual proteins. Therefore, to compare the two methods, we depleted several proteins in HeLa cells by RNAi. We then analyzed the same extracts from control and KD cells by the Odyssey and by MS Western and checked if they reported concordant changes in protein abundances.

For MS Western quantification we designed a 72 kDa chimera protein CP03 comprising 47 proteotypic peptides from six human proteins: CAT, TUBA, PLK1, AKT1, GAPDH and PTEN (Supplementary Fig. S5, S6 and Supplementary Table S3) for each of which commercial monoclonal antibodies are available. Both Odyssey and MS Western determined quantitatively concordant levels of endogenous TUB, PLK1, CAT and AKT1 in KD-cells as compared to the luciferase control (Supplementary Fig. S16). Interestingly, MS Western could separately quantify the relative abundances of TUBA4A and TUBA1A sharing 96% of the full-length sequence identity. The abundance of both proteins was affected by RNAi at the same extent, however they were not distinguished by the Odyssey (Supplementary Fig. S16F).

We used the same proteins to benchmark the dynamic range and sensitivity of MS Western quantification in comparison to the Odyssey. To this end, samples of successively diluted total protein extract from HeLa cells were loaded on two different gels. We adjusted the loaded volumes such that, for each dilution, the same amount of protein extract (here exemplified as an equivalent number of extracted cells) was subjected to Odyssey imaging and injected into the LC-MS/MS. Therefore, protein molar amounts (as determined by MS Western) and integrated intensities of protein bands (determined by Odyssey) could be correlated with no further adjustment (Fig. 5 B, C).

Fig. 5D and 5E compare the dilution plots of TUBA quantified by MS Western (as TUBA1A) and by the Odyssey (as TUBA1A and TUBA4A), respectively. In WB experiments GAPDH served as a loading control and its abundance followed the dilution of HeLa extract. In MS Western an equal amount of CP03 was co-digested with each sample and therefore its constant abundance evidenced the analyses consistency. Compared to the Odyssey, MS Western showed at least 60-fold better sensitivity (down to 1.5 fmol of TUBA1A) along with higher dynamic range (>16000-fold) and excellent linearity (*r*^2^ = 0.9983) (Fig.5E). In the middle of the plot (Fig.5D) the Odyssey also showed good linearity towards both GAPDH and TUB1A/4A. However, at lower loadings the Odyssey did not recognize the target proteins, while at higher loadings the system incorrectly integrated the abundance of irregular-shaped protein bands and lost the response linearity. Expectantly, we could also quantify proteins from gel lanes with no visible protein bands (Fig.5C, E); for better visualization we also present the expanded gel image showing the four most diluted samples corresponding calibration plots (Supplementary Fig. S17).

Contrary to WB that is often pestered with false positives, MS Western seems to be more prone to false negatives, particularly if overwhelming background abolishes the ionization of proteotypic peptides and / or if target proteins are extremely low abundant. Nevertheless, we were able to quantify proteins at the sub-femtomole to hundred attomole range in complex biological extracts. Supplementary Fig.S18 presents calibration plots for sub-femtomole quantification of BSA spiked into a total protein extract from *E.coli*. Supplementary Fig. S19 presents the attomole quantification of 229 kDa transmembrane protein axotactin in two biological conditions from a total extract of *Drosophila* eye using 264 kDa chimera CP04 (Supplementary Fig. S7, S8 and Supplementary Table S4). The figure provides BPC and XIC traces as well as FT MS spectra of light and heavy forms of the two axotactin peptides and of five reference peptides from BSA acquired in the same quantification experiment. We also provide the ratios of relative abundances of light and heavy peptides suggesting that even at the hundred attomoles level protein quantification with both peptides were concordant.

We note that MS Western workflow does not enhance the detection sensitivity compared to conventional GeLC-MS/MS and might fail to quantify very low abundant proteins that might still be detectable by classical Western. Although in MS Western a linear calibration plot could be extended down to 100-attomoles of a gel separated protein standard (Supplementary Fig. S20), it is nevertheless advisable to always start with preliminary GeLC-MS/MS experiments before designing CPs to make sure that target proteins are detectable and to select the optimal constellation of proteotypic peptides.

### Absolute quantification of histones in zebrafish embryos

Histones make up the basic unit of chromatin, the nucleosome. Each nucleosome consists of 147 base pairs of DNA wrapped around a histone octamer comprising two copies of each of the four core histones: H2A, H2B, H3, and H4. Embryos inherit histones from the mother (55) and previous WB analyses suggested that, despite embryo growth, histones level is stable during the early stages of development in both *Xenopus* (56) and zebrafish (57), but increases rapidly upon genome activation. We reasoned that predictable stoichiometry and time course of their total content make embryonic histones a good model for validating MS Western in *in vivo* experiments.

We employed MS Western to determine the molar content of H2A, H2B, H3 and H4 in zebrafish embryos at ten developmental stages ranging from 1-cell stage to Shield stage. Zebrafish embryos (1-cell stage *n*=5; otherwise *n*=10) were collected and (except for 1-cell stage) manually deyolked (58). Proteins extracted from deyolked embryos were subjected to SDS PAGE. Slabs corresponding to the Mr range of *ca*. 5 to 25 kDa that contained all core histones were excised from each gel and histones were quantified using the bands of CP02 and BSA. We observed that ratios of relative abundances of light and heavy forms of proteotypic peptides (Fig. 6A) varied by less than 10% in all embryogenesis stages and concluded that endogenous peptides were not harbouring PTMs. Accurate (CV <10%) absolute quantification of four histones (Fig 6B; Supplementary Table S9) revealed that, in contrast to previous reports, histones content steadily increased already during the early stages of development (59). This suggested that in zebrafish embryos mothers deposit both histone proteins and corresponding mRNA that are translated before the onset of zygotic transcription around the 1000-cell stage. As expected, after genome activation, histones content increased more rapidly.

**Fig. 6.**
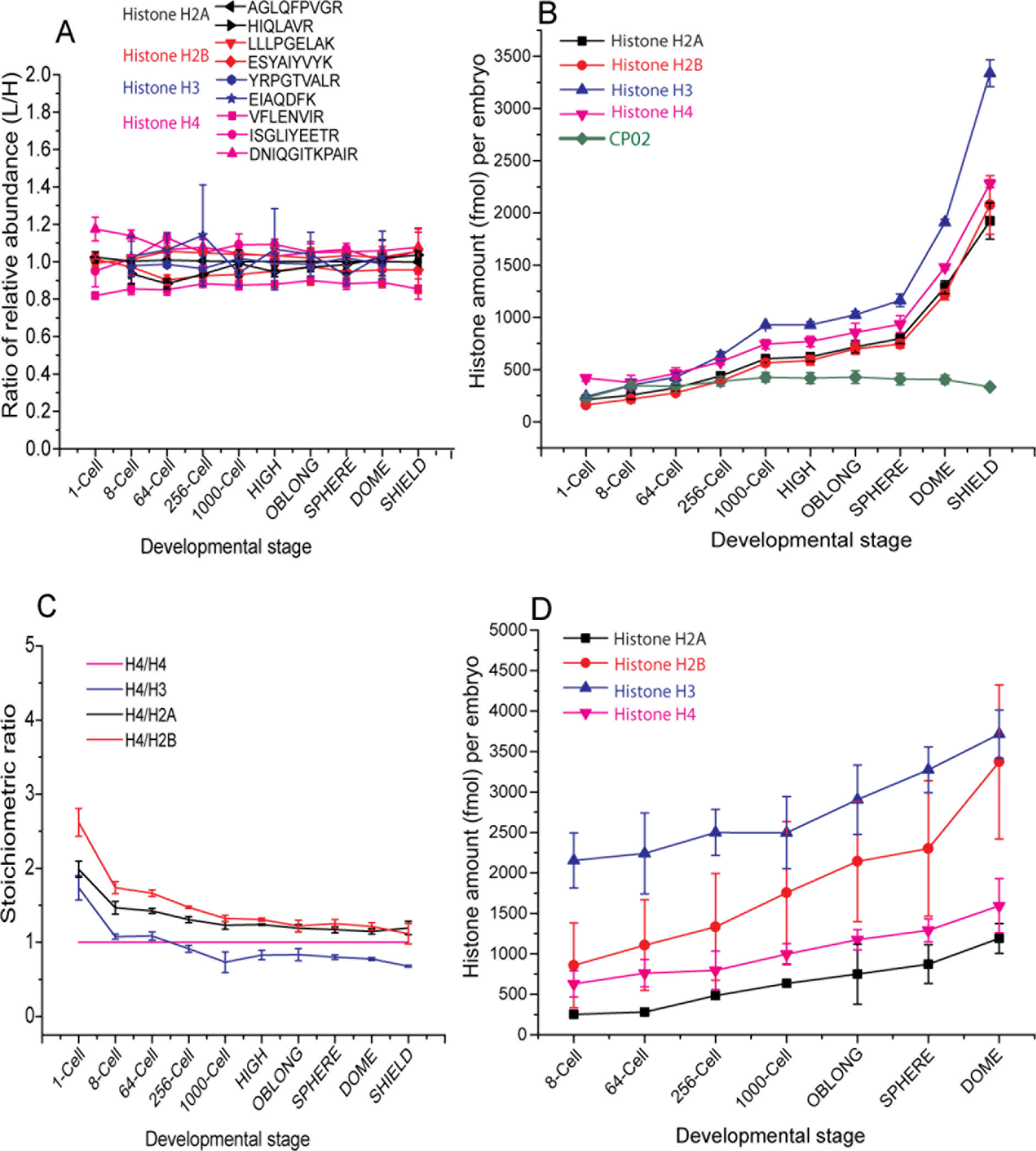
Absolute quantification of core histones in zebrafish embryos. (**A**) Ratios of relative abundances of light and heavy proteotypic peptides produced from endogenous histones at different developmental stages and from CP02, respectively. (**B**) Molar content of individual histones per zebrafish embryo as quantified by MS Western. (**C**) Stoichiometric ratios between the four histones at as determined by MS Western. (**D**) Histones content quantified by LI-COR Odyssey. Data presented as mean ±SD (*n*=3).

Molar quantities of individual histones were consistent with the expected equimolar stoichiometry (Fig. 6C). At the very early stages of development the histones stoichiometry slightly, yet consistently deviated from equimolar ratios, however that were closely approached at the later stages. We note that MS Western quantification only reflected the total content of histones, irrespectively if they were assembled into a nucleosome or associated with chaperones (55, 60).

We also quantified histones by the Odyssey (Fig. 6D) and compared it to MS Western (Supplementary Fig. S21). We found that the quantification by Odyssey was generally inconclusive and stoichiometry between individual histones was nowhere close to the expected equimolar ratios. This, once again, highlighted that WB quantification is extremely sensitive towards the quality of antibodies and protein standards.

Taken together, MS Western enabled precise and consistent quantification of the molar amounts of four core histones at the sub-picomole per embryo level directly from total protein extracts and revealed their unexpected individual dynamics during early embryogenesis.

## CONCLUSIONS AND PERSPECTIVES

We argue that MS Western is a practical and technically simple solution for the accurate multiplex targeted absolute quantification of proteins. In MS Western workflow SDS PAGE circumvents the limited solubility of both target and chimera proteins; it also removes interfering buffers and detergents, including SDS. Pre-separation of total protein extracts improves the dynamic range and sensitivity of quantification. Protein quantification relies on multiple proteotypic peptides and the concordance between the relative abundances of matching pairs of heavy and light peptides provides independent validation of the quantification consistency. However, one conceptual limitation of MS Western is that it is difficult to predict if a particular low abundant protein (*e.g*. a transcription factor) could be detectable by GeLC-MS/MS. In our experience the detection of proteins by classic Western blotting or projecting their abundances from transcriptomics did not substitute preliminary analyses by GeLC-MS/MS and it is advisable to establish the scope of detectable proteins before planning systematic MS Western experiments. However, once target proteins have been detected, their accurate absolute multiplexed quantification becomes a straightforward technical task.

In the future it would be interesting to explore if a single, reasonably large CP could cover all essential members of metabolic and/ or signalling pathways and allow us to relate their molar abundances to molar concentrations of corresponding metabolites. It would also be intriguing to explore if CP could be used as templates to incorporate site-specific post-translational modifications by some chemical or genetic means.

## AUTHOR CONTRIBUTIONS

MK, MG and AS designed the study. MK, AB, and DD designed and produced protein chimeras. MK performed MS Western benchmarking and validation experiments and quantified zebrafish histones. SRJ and NLV performed experiments with zebrafish embryos. MA and FB performed KD experiments in HeLa cells. MK and AS wrote the draft that was jointly revised by all co-authors.

## ACKNOWLEDGEMENTS

Work in AS and NLV laboratories was supported by MPG core funding. AS laboratory was supported by KFO249 and TRR83 (Project A17) grants from Deutsche Forschungsgemeinschaft; NLV laboratory by Human Frontier Science Program Career Development Award (CDA00060/2012). Work in the FB laboratory was supported by the Excellence Initiative of the German Federal and State Governments (Institutional Strategy, measure ‘support the best ZUK 64’). MK and SRJ are recipients of IMPRS PhD student fellowship. The authors are grateful for the members of AS, NV and FB laboratories for useful discussion and expert technical support. We are particularly grateful for Dr. Henrik Thomas (MPI CBG) for developing useful data processing tools.

